# Ciliary and rhabdomeric photoreceptor-cell circuits form a spectral depth gauge in marine zooplankton

**DOI:** 10.1101/251686

**Authors:** Csaba Verasztó, Martin Gühmann, Huiyong Jia, Vinoth Babu Veedin Rajan, Luis A. Bezares-Calderón, Cristina Piñeiro Lopez, Nadine Randel, Réza Shahidi, Nico K. Michiels, Shozo Yokoyama, Kristin Tessmar-Raible, Gáspár Jékely

**Author notes:** Present address: Janelia Research Campus, USA. Present address: EMBL Heidelberg Meyerhofstraße 1, 69117 Heidelberg, Germany. These authors contributed equally to this work (Csaba Verasztó, Martin Gühmann).

## Abstract

Ciliary and rhabdomeric photoreceptor cells represent two main lines of photoreceptor evolution in animals. The two photoreceptor-cell types coexist in some animals, however how they functionally integrate is unknown. We used connectomics to map synaptic paths between ciliary and rhabdomeric photoreceptors in the planktonic larva of the annelid *Platynereis* and found that ciliary photoreceptors are presynaptic to the rhabdomeric circuit. The behaviors mediated by the ciliary and rhabdomeric cells also interact hierarchically. The ciliary photoreceptors are UV-sensitive and mediate downward swimming to non-directional UV light, a behavior absent in ciliary-opsin knockouts. UV avoidance antagonizes positive phototaxis mediated by the rhabdomeric eyes so that vertical swimming direction is determined by the ratio of blue/UV light. Since this ratio increases with depth, *Platynereis* larvae may use it as a depth gauge during planktonic migration. Our results revealed a functional integration of ciliary and rhabdomeric photoreceptors with implications for eye and photoreceptor evolution.

## Introduction

Comparative studies have identified two major photoreceptor cell-types in bilaterians, the rhabdomeric‐ and the ciliary-type photoreceptor cells (rPRC and cPRC, respectively) (Arendt, 2003; Arendt et al., 2004; Eakin, 1979; Erclik et al., 2009). These cells have distinct morphologies and express different classes of opsins. Rhabdomeric PRCs with microvillar specializations are the visual photoreceptors in most protostome eyes and also exist in pigmented eyespots in both protostomes and deuterostomes (Arendt and Wittbrodt, 2001; Braun et al., 2015; Nakao, 1964). Vertebrate visual eyes have cPRCs with ciliary membrane specializations. In invertebrates, cPRCs can occur in pigmented eyespots, in simple eyes, or as brain photoreceptors not associated with pigment cells (Arendt et al., 2004; Barber et al., 1967; Randel and Jékely, 2016; von Döhren and Bartolomaeus, 2018). Opsin expression generally correlates with the morphological type with cPRCs expressing copsins and rPRCs expressing r-opsins (Arendt, 2003; Arendt et al., 2004; Randel et al., 2013; Vopalensky et al., 2012) with occasional coexpression (Davies et al., 2011; Vöcking et al., 2017). Given their broad distribution, both photoreceptor-cell types likely coexisted in the ancestral bilaterian. The two types still coexist in several marine animals, including cephalochordates, some mollusks, flatworms, and annelids (Arendt et al., 2004; McReynolds and Gorman, 1970; Randel and Jékely, 2016; Vopalensky et al., 2012) and form parts of pigmented or non-pigmented light sensitive structures. Understanding how the two photoreceptor-cell types integrate at the functional and circuit levels in these organisms could clarify the history of eyes and photoreceptor cells.

Here we study the planktonic larva of *Platynereis dumerilii*, a marine annelid that has both photoreceptor cell types. In *Platynereis*, non-pigmented brain cPRCs with ramified cilia express a ciliary type opsin (c-opsin1) (Arendt et al., 2004) and coexist with r-opsin-expressing rPRCs that are part of the pigmented visual eyes (adult eyes) and eyespots (Arendt et al., 2002; Jékely et al., 2008; Randel et al., 2013; Randel et al., 2014). The pigmented eyes mediate early‐ and late-stage larval phototaxis (Gühmann et al., 2015; Jékely et al., 2008; Randel et al., 2014). The cPRCs in *Platynereis* have been proposed to regulate melatonin production and to entrain the circadian clock to ambient UV light (Arendt et al., 2004; Tosches et al., 2014; Tsukamoto et al., 2017). However, the function of the cPRCs in *Platynereis* and how they interact with rPRCs is still unknown.

## Results

To identify synaptic connections between the cPRC and rPRC circuits, we used a serial-section transmission electron microscopy (ssTEM) dataset spanning the entire body of a 72 hours post fertilization (hpf) *Platynereis* larva (Randel et al., 2015). Previously, we reported the synapse-level connectome of the rPRCs from the visual eyes and eyespots, (Randel et al., 2014; Randel et al., 2015) and the direct postsynaptic circuit of the four cPRCs with ramified cilia (Williams et al., 2017). Here, we analyzed all synaptic connections between the cPRC and rPRC circuits (Figure 1) based on the reconstruction of all neurons and their synapses in the larval head (598 sensory, 616 inter-, 18 motorneurons; unpublished data). We identified three sites of contact. First, the RGW interneurons (IN^RGW^), which are directly postsynaptic to the cPRCs (Williams et al., 2017), synapse on the Schnörkel interneurons (IN^sn^), which represent the premotor interneurons in the visual circuit (Figure 1E-G and Movie 1)(Randel et al., 2014). Second, we identified six interneurons (IN^preMN^) that are postsynaptic to the RGW neurons and presynaptic to the ventral motorneurons (vMNs), the motorneurons of the visual circuit (Figure 1EG)(Randel et al., 2014). Third, two putative mechanosensory neurons in the median head (MS1 and MS2) are postsynaptic to the RGW cells and presynaptic to the vMNs and IN^pro^ interneurons of the visual circuit (Figure 1E-G and Movie 1). We did not find any neurons that were postsynaptic to the rPRC circuit and presynaptic to the cPRC circuit. Thus, the cPRC circuit feeds hierarchically into the visual rPRC circuit (Figure 1G). This suggests that the cPRCs could influence phototaxis, a behavior mediated by the rhabdomeric eyes and eyespots (Gühmann et al., 2015; Jékely et al., 2008; Randel et al., 2014).

**Figure 1.**
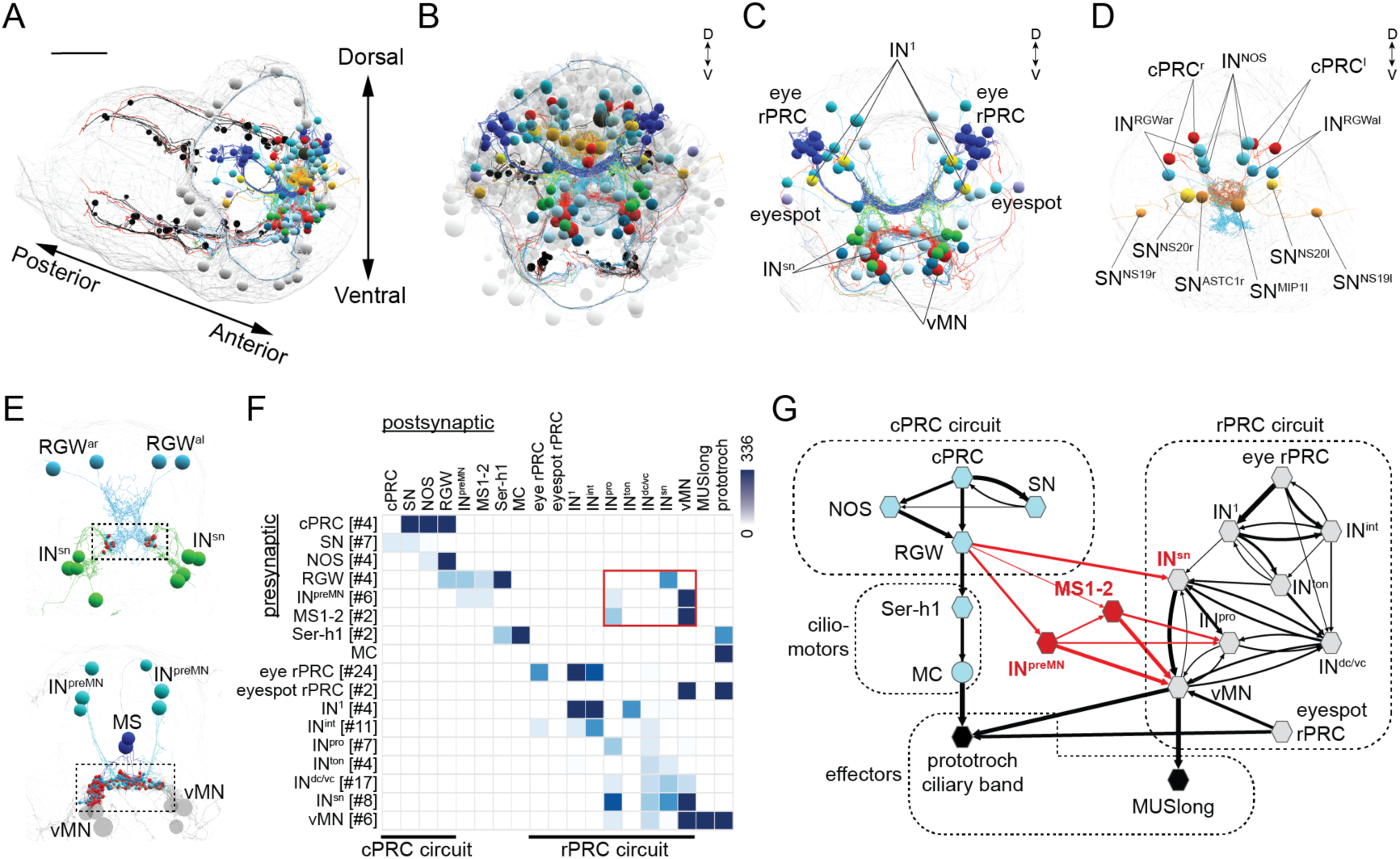
Wiring diagram of cPRC and rPRC circuits in the *Platynereis* larval head. (A) Circuits were reconstructed from a whole-body ssTEM volume of a 72 hpf larva. (B) All cells of the cPRC and rPRC circuits in the larval head (in color). All other neurons are shown in grey. (C) All neurons of the rPRC circuit. (D) The four cPRCs and all neurons directly postsynaptic to them. (E) Connections between the cPRC and rPRC circuits. Top panel: the four RGW neurons are presynaptic to the IN^sn^ cells. Synapses from RGW to IN^sn^ are in red (boxed area). Bottom panel: the six IN^preMN^ cells and the two MS cells are presynaptic to the ventral motorneurons. (F) Grouped connectivity matrix of the cPRC and rPRC circuits. The connections from the cPRC to the rPRC circuit are boxed. (G) Wiring diagram of the cPRC and rPRC circuits. Synaptic connections from the cPRC to the rPRC circuit are in red.

What is the role of cPRCs in larval behavior and how do they influence phototaxis? First, to estimate the sensitivity of the cPRCs, we used ssTEM to reconstruct their morphology and to measure their total sensory membrane surface area (Figure 2A-D). The area was 276 μm^2^, which is approximately 10 times smaller than the total disk membrane surface area of rat rods (Mayhew and Astle, 1997). This suggests that *Platynereis* cPRCs are sensitive enough to mediate acute light sensation. Next, we expressed *Platynereis* c-opsin1 in COS1 cells, reconstituted it with 11-cis-retinal and purified it (Yokoyama, 2000). The reconstituted pigment absorbed in the UV range with a λ-max of 384 nm in the dark spectrum and a λ-max of 370 nm in the dark-light difference spectrum (Figure 2E), in agreement with a recent report (Tsukamoto et al., 2017).

**Figure 2.**
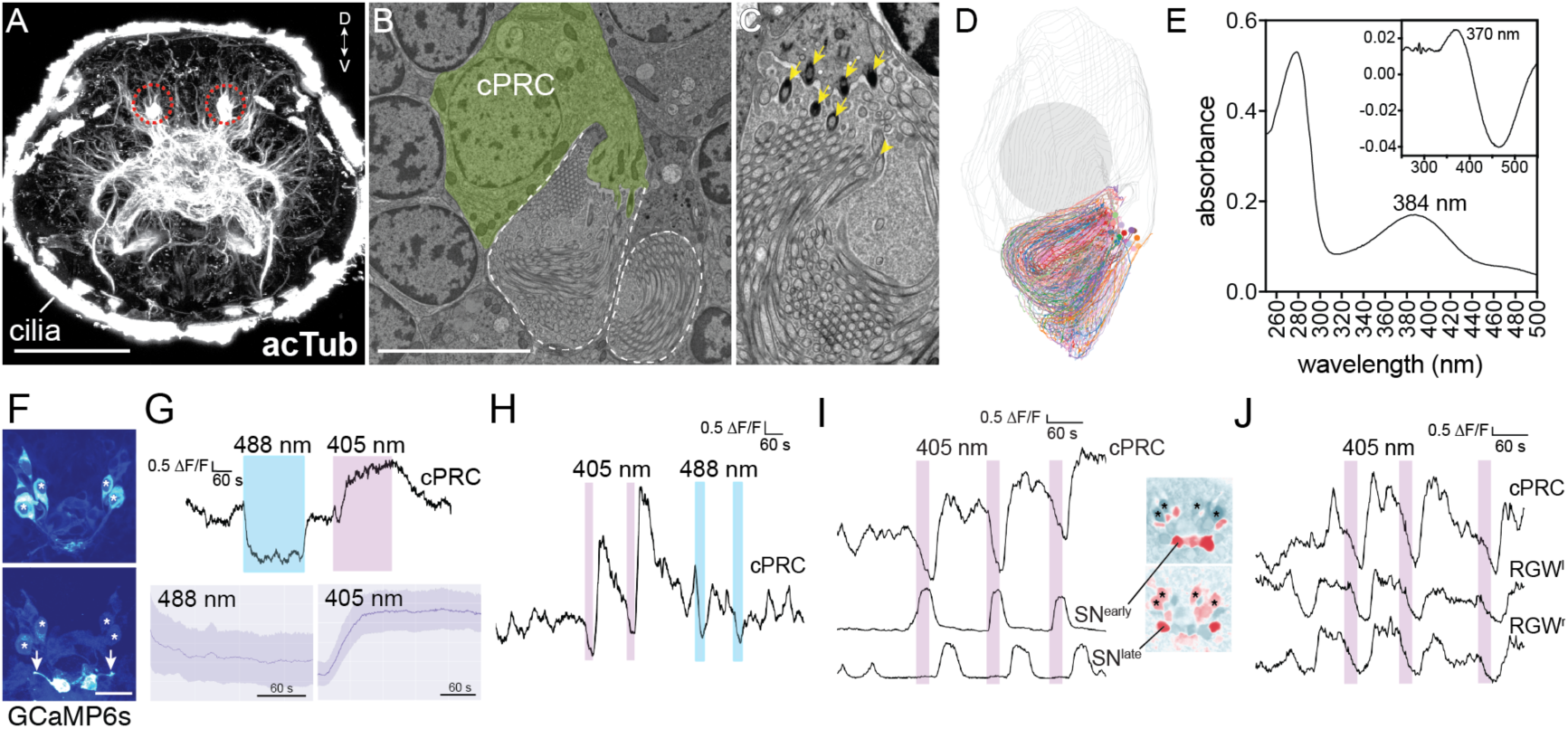
Light responses of brain ciliary photoreceptors and their downstream circuitry in *Platynereis* larvae. (A) Acetylated tubulin staining of a 72 hpf larva. Ramified sensory cilia of cPRCs are marked with circles. (B) TEM image of cPRCs. (C) TEM image of a cPRC with sensory cilia. Arrows mark basal bodies. (D) Serial TEM Reconstruction of the sensory cilia of a cPRC. (E) Absorption spectrum of purified *Platynereis* c-opsin1. Inset shows the dark-light difference spectrum. (F) Top: high GCaMP6s signal in cPRCs during imaging conditions. Asterisks mark cPRC nuclei. Bottom: Activation of two sensory neurons upon UV stimulation of cPRCs. (G) Top: representative example of cPRC response to prolonged local 488 nm and 405 nm stimulation. Colored boxes show duration of stimulation. Bottom: average cPRC response during continuous 488 nm and 405 nm stimulation. Data show mean and s.d. of mean, 405 nm N>30, 488 nm N=8. (H) Responses of a cPRC to repeated 405 nm and 488 nm (duration: 20 sec) stimulation. (I) Responses of SN^early^ and SN^late^ sensory neurons to cPRC UV stimulation. Correlation images are shown for SN^early^ and SN^late^. Asterisks mark cPRC nuclei. (J) Responses of RGW cells to UV stimulation of a cPRC. Scale bars: (A) 50 μm (B) 10 μm, (F) 20 μm.

To investigate how cPRCs respond to light, we did calcium imaging in larvae ubiquitously expressing the calcium sensor GCaMP6s (Chen et al., 2013; Verasztó et al., 2017). When imaged with a low-intensity 488 nm laser, the cPRCs had a high resting calcium level and the GCaMP6s signal highlighted their sensory cilia. We could thus identify the four cPRCs without stimulation (Figure 2F). When the cPRC cilia were stimulated locally (Supplementary Figure 1) for 5 min with 405 nm light (140-250 times more photons on the sensory cilia than by the imaging laser), the calcium level dropped transiently at the cPRC somata and then increased strongly. This indicates transient cPRC hyperpolarization followed by depolarization. However, when the cPRCs were stimulated with a 488 nm laser of equal photon flux, they showed only prolonged hyperpolarization but no depolarization (Figure 2G). When the 405 nm stimulation was switched off during the hyperpolarization phase (after 20 sec), the cPRCs still depolarized afterwards (Figure 2H).

405 nm stimulation of the cPRCs also changed the activity of other neurons in the larval brain. Four neurons followed the activity pattern of the cPRCs (Figure 2J). These cells correspond by position to the four RGW interneurons, which together with four NOS interneurons represent the direct interneuron targets of the cPRCs (Figure 1G)(Williams et al., 2017). Additionally, two flask-shaped sensory neurons (SN^early^) in the middle of the anterior nervous system depolarized upon stimulus onset. Two further sensory cells (SN^late^) flanking the SN^early^ cells depolarized later, in parallel with the rising phase of cPRC activity (Figure 2I). These four cells correspond by position to four sensory-neurosecretory neurons that are postsynaptic to the cPRCs in the anterior nervous system (SN^MIP1^, SN^NS20^)(Williams et al., 2017). We focused on the responses of the four SN neurons to repeated 405 and 488 nm stimulation of the cPRCs. While 405 nm light induced depolarization, the SN neurons did not respond upon 488 nm stimulation of the cPRCs (Supplementary Figure 2). Thus, postsynaptic neurons are activated by violet but not blue stimulation of the cPRCs.

To characterize how *Platynereis* larvae react to UV light, we assayed larval swimming behavior in a vertical column setup. Since *Platynereis* larvae are strongly phototactic to a broad spectrum of light (Gühmann et al., 2015; Jékely et al., 2008), we illuminated the setup equally from two opposite sides with non-directional UV light so that the larvae could not respond with directional phototaxis (Figure 3A).

**Figure 3.**
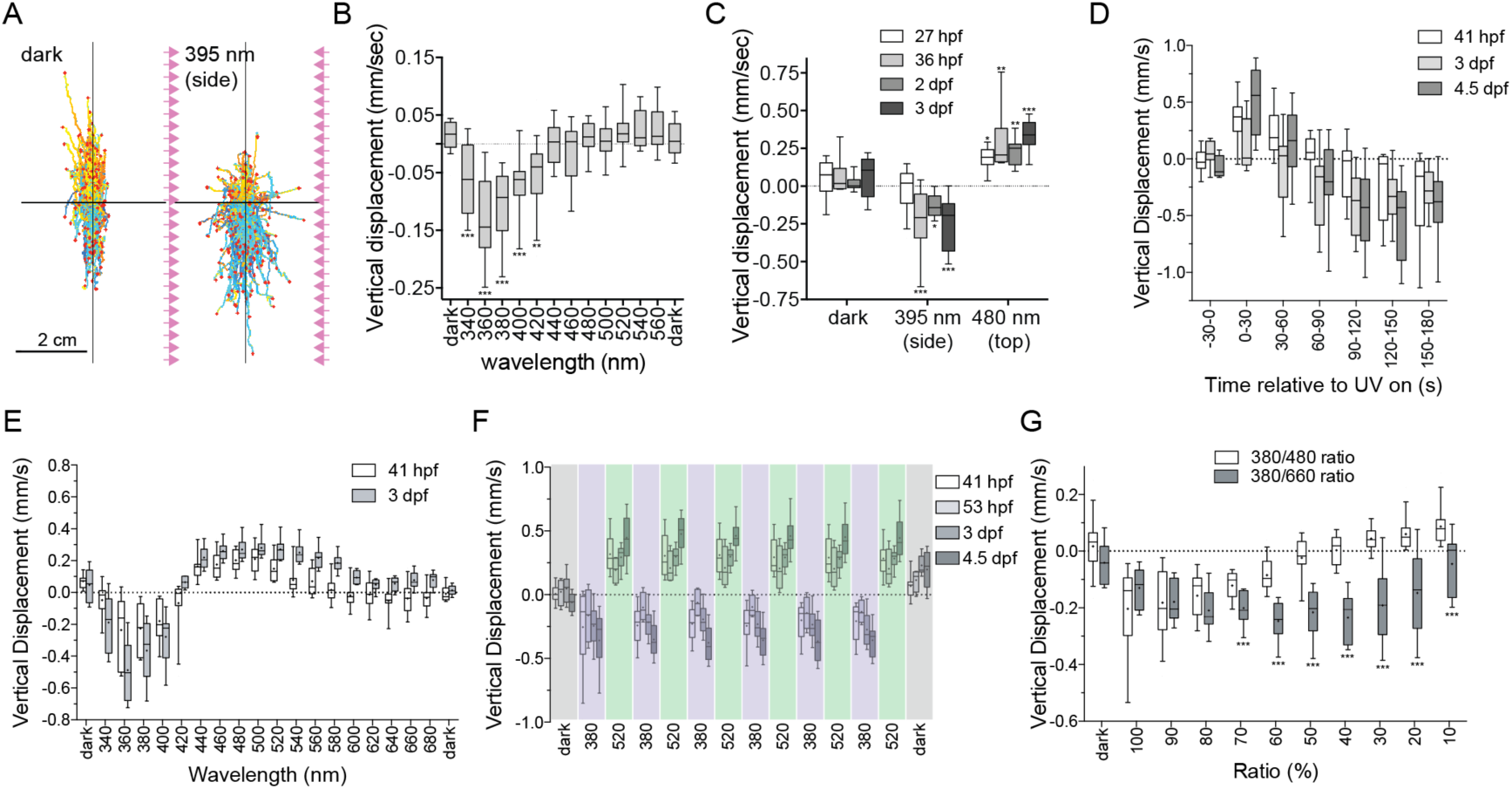
UV avoidance and phototaxis form a ratio-chromatic depth gauge in *Platynereis* larvae. (A) Larval trajectories recorded in a vertical column in the dark (left) and under illumination with UV light from the side (right). (B) Action spectrum of non-directional light avoidance in 48 hpf larvae. (C) Developmental onset of UV-avoidance behavior and phototaxis in *Platynereis* larvae. (D) Time course (30 sec bins) of vertical swimming in different larval stages following 380 nm illumination from above. (E) Action spectrum of vertical swimming in early‐ and late-stage larvae, under stimulus light coming from the top of the column. (F) Repeated switching between upward and downward swimming in different larval stages under 380 and 520 nm stimulus light coming from above. (G) Ratio-dependent switch in vertical swimming direction in 3-day-old larvae stimulated from above with different mixtures of monochromatic light. T-tests with Holm-Sidak correction (alpha=0.05) were used. Significant differences are indicated (*** p-value < 0.005).

When the larvae were stimulated with UV light, they started to swim downward. To characterize the wavelength dependence of this behavior, we used monochromatic stimulation of different wavelengths provided from two sides of the setup. The larvae swam down to UV-violet light (between 340-420 nm) but not to longer wavelengths (>420 nm, Figure 3B). This action spectrum closely matches the absorption spectrum of c-opsin1 (Figure 2E).

During the course of larval development, the appearance of UV avoidance approximately coincides with the morphological differentiation of cPRCs (after 32 hpf, as judged by the appearance of the ramified cilia)(Figure 3C), but not with the differentiation of the eyespots (at 24 hpf), or adult eyes (at 72 hpf)(Fischer et al., 2010; Jékely et al., 2008; Randel et al., 2014). Thus, *Platynereis* larvae show UV-light avoidance that is independent of phototaxis and is likely mediated by the UV/violet-responding non-pigmented cPRCs.

To study how UV avoidance interacts with rhabdomeric-eye-mediated phototaxis, we stimulated the larvae in the vertical column with directional 380 nm light from above. Both early‐ (41 hpf) and late-stage (3 and 4.5 dpf) larvae swam first upward, towards the light, and then swam downward (Figure 3D, Movie 2). This upward phase was not observed when the larvae were illuminated uniformly from the side (data not shown). This suggests that the upward-swimming phase is phototaxis, which is then overwritten by the UV-avoidance response.

Next, we measured how the larvae responded to different wavelengths of light coming from the top of the vertical column. In response to illumination with 340-400 nm light, larvae swam downward. In response to illumination with 440-540 nm light (early-stage) or 440-600 nm light (late-stage), larvae swam upward (Figure 3E). The swimming direction could be switched several times by changing the wavelength of the illuminating light source from 380 nm to 520 nm (Figure 3F), demonstrating that this behavioral switch does not habituate even after sustained exposure. This switching in behavioral response cannot be explained as a wavelength-dependent alternation between positive and negative phototaxis, since early-stage larvae are exclusively positively phototactic (Jékely et al., 2008)(until 72 hpf (Randel et al., 2014)), yet already show the behavioral switch (Figure 3E, F).

Since blue and UV/violet light attenuate differently in seawater, the ratio of blue to UV/violet light increases with depth (Gühmann et al., 2015). Strong UV/violet light at the ocean’s surface is expected to induce larvae to swim downward as an avoidance response, and relatively strong blue light in deeper waters is expected to induce phototactic upward swimming. Such depth-dependent behavioral switching could form a ratio-chromatic depth-gauge (Nilsson, 2009). To test this, we exposed larvae to mixed wavelength light containing different photon ratios of UV (380 nm) to blue (480 nm) light. At high 380/480 ratios, larvae swam downward, while at low ratios larvae swam upward. At a 40% 380/480 ratio, larvae remained at a constant average depth (Figure 3G). Mixing UV light with 660 nm red light (a wavelength to which larvae do not respond phototactically at the intensity used (Figure 3E)) did not induce upward swimming at any 380/660 ratio (Figure 3G).

To test whether c-opsin1 mediates the UV response in *Platynereis* larvae, we used a *c-opsin1 Platynereis* knockout line generated by TALEN-mediated genome editing (Rajan VB et al. submitted). The TALENs targeted the third exon of *c-opsin1* and induced an 8 bp deletion (Figure 4A). In homozygous *c-opsin1*^*Δ8/Δ8*^ larvae, the cPRCs had low resting calcium level and neither hyperpolarized nor depolarized upon UV stimulation (Figure 4B, C). Thus, c-opsin1 in the *Platynereis* cPRCs is required for an elevated resting calcium level in the dark state and for hyperpolarization and subsequent depolarization upon UV exposure. The high resting calcium level in cPRCs is indicative of a dark current, a characteristic of vertebrate rods and cones (Hagins et al., 1970).

**Figure 4.**
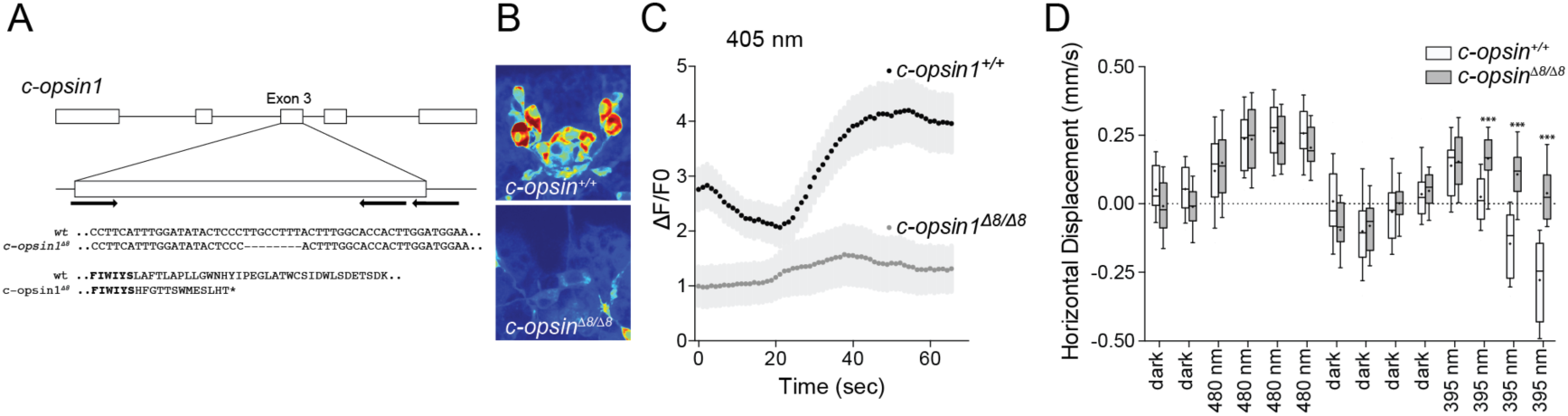
*c-opsin1* knockout larvae lack UV/violet responses. (A) Schematic of the *Platynereis c-opsin1* gene and the *c-opsin1*^*Δ8/Δ8*^ mutation. (B) Background GCaMP6s signal during calcium imaging in wild type and *c-opsin1*^*Δ8/Δ8*^ mutant larvae. (C) Calcium responses to 70 sec 405 nm stimulation in wild type and mutant larvae. (D) Vertical swimming in wild type and mutant larvae stimulated with blue (480 nm) and UV (395 nm) light from above. P-values: ***<0.001.

Next, we tested behavioral responses to light in *c-opsin1*^*Δ8*^ mutant larvae. Similar to wild type larvae, homozygous *c-opsin1*^*Δ8/Δ8*^ larvae swim upward (positive phototaxis) in response to 480 nm light. However, *c-opsin1*^*Δ8/Δ8*^ larvae have a defective UV-avoidance response and continue to swim upward, showing positive phototaxis in response to UV light, whereas wild type larvae swim downward (Figure 4D). These results show that c-opsin1 mediates UV-avoidance and is required for the depth gauge. If c-opsin1 is missing, UV light induces the default positive phototactic behavior. In wild-type larvae, UV avoidance overrides positive phototaxis after 30 sec (Figure 3D) and the two opposing behaviors cancel each other out at a set UV/blue ratio (Figure 3G). Under blue illumination, the rhabdomeric eyes mediated phototaxis, and the cPRC are inhibited (Figure 2G).

## Discussion

Our results are consistent with the presence of a ratio-chromatic depth-gauge in the planktonic larvae of *Platynereis dumerilii*. The depth gauge is formed by the antagonistic interaction of two distinct behaviors, mediated by two distinct types of photoreceptor systems.

We found that the cPRC circuit connects to the circuit of the rhabdomeric eyes via distinct synaptic pathways, providing inputs at the level of the IN^sn^, IN^pro^ and the vMN cells. The exact mechanism of UV avoidance and how this behavior overrides phototaxis at the neuronal level, remain to be elucidated. Since the cPRCs and their direct postsynaptic targets are strongly neuroendocrine (Williams et al., 2017), it is possible that neuroendocrine volume transmission also contributes to the behavioral antagonism, along with the synaptic connections that we identified in the wiring diagram.

One outstanding question in eye evolution is why did the ciliary and rhabdomeric PRC types evolve (Nilsson, 2009; Nilsson, 2013). Our findings suggest that the two types may have evolved antagonistic functions early in evolution. Invertebrates then elaborated on the rhabdomeric, and vertebrates on the ciliary type (Arendt et al., 2004). According to one hypothesis, as the brain cPRCs were recruited for vision, the rPRCs evolved into the retinal ganglion cells (Arendt, 2003; Arendt, 2008). This scenario is supported by the expression of melanopsin, an r-opsin ortholog, in the intrinsically photosensitive retinal ganglion cells (Hattar et al., 2002; Hattar et al., 2003; Koyanagi et al., 2005; Lucas et al., 2001; Panda et al., 2003). Although we need more comparative data to test this model, we hypothesize that the cell-type mosaic of the vertebrate retina may have originated from the hierarchical integration of distinct cPRC and rPRC circuits mediating antagonistic behaviors, as observed in *Platynereis* larvae.

## Methods

### *Platynereis dumerilii* culture

Larvae of *Platynereis dumerilii* were cultured at 18°C in a 16 h light 8 h dark cycle until experiments. Larvae were raised to sexual maturity according to established breeding procedures.

### Estimation of cPRC sensory membrane surface

We used serial-sectioning transmission electron microscopy to analyze cPRC sensory morphology. Each cPRC has 15 basal bodies, each basal body gives rise to one cilium that branches at its base to 3-9 branches, each being supported by at least one microtubule doublet. The branches have an average diameter of 130 nm (st.dev. 19 nm) and a total length of 677 μm (measured in one cPRC). This represents a total membrane surface area of approximately 276 μm^2^.

### *In vitro* absorption spectrum of c-opsin1

Recombinant c-opsin1 was purified and analyzed following (Yokoyama, 2000). Full-length *Platynereis c-opsin1* was amplified with primers that introduced EcoRI, Kozak and SalI sequences, and cloned into the expression plasmid pMT5. C-opsin1 was expressed in COS1 cells and incubated with 11-cis-retinal (a gift from Dr. Rosalie K. Crouch at Storm Eye Institute) to regenerate the photopigment. The pigment was purified with immobilized 1D4 (The Culture Center, Minneapolis, MN) in buffer W1 (50 mM N-(2-hydroxyethyl) piperazine-N’-2-ethanesulfonic acid (HEPES) (pH 6.6), 140 mM NaCl, 3mM MgCl2, 20% (w/v) glycerol and 0.1% dodecyl maltoside). The UV/VIS spectrum of the pigment was recorded at 20°C with a Hitachi U-3000 dual beam spectrophotometer. The pigment was bleached for 3 min with a 60 W standard light bulb equipped with a Kodak Wratten #3 filter at a distance of 20 cm.

### *c-opsin1* sequencing and genotyping

For genotyping of the *c-opsin1* locus, genomic DNA was isolated from single larvae, groups of 6-20 larvae, or from the tails of adult worms. The DNA was amplified by PCR (primers: 5’-AGCCTTCATTTGGATATACTCCC-3’, 5’-TTATAAACGATGGAACTTACCTTTCTG-3’) with the dilution protocol of the Phusion Human Specimen Direct PCR Kit (Thermo Scientific). The PCR product was sequenced directly with a nested sequencing primer (5’-ATGAGACCATACGAAACCAC-3’). A mixture of wild-type and deletion alleles in a sample gave double peaks in the sequencing chromatograms, with the relative height of the double peaks reflecting the relative allele ratio in the sample.

### Photostimulation

We used a variety of light sources for photostimulation, depending on the experimental setup. In the calcium imaging experiments we used the available laser lines (405 and 488 nm; Showa Optronics, Tokyo) operating in continuous mode. The power of the lasers was measured with a microscope slide power sensor (S170C; Thorlabs, Newton, USA). The typical power for stimulation was 5.59 μwatts for the 405 nm laser and 4.62 μwatts for the 488 nm laser. These values correspond to 1.1e+13 photons/sec. For behavioral assays, we used either UV LEDs (395 nm peak wavelength) or a monochromator (Polychrome II, Till Photonics) for which we quantified photon irradiances across the spectrum (3-4x10^18^ photons/sec/m^2^) (Gühmann et al., 2015). We refer to color according to these wavelength ranges: UV < 400 nm, violet 400-450 nm, blue 450-490 nm (450-460 royal blue), cyan 490-520 nm.

### Calcium imaging

For calcium imaging, 36-52 hpf larvae were used. Experiments were conducted at room temperature in filtered natural seawater. Larvae were immobilized between a slide and a coverslip spaced with adhesive tape. GCaMP6s mRNA (1 mg/μl) was injected into zygotes as described previously (Randel et al., 2014). Imaging was done on an Olympus Fluoview-1200 (with a UPLSAPO 60X water-immersion objective, NA 1.2) or on a Leica TCS SP8 (with a 40X water-immersion objective, NA 1.1) microscope with a frame rate of 1.25/sec and an image size of 254x254 pixels. The larvae were stimulated in a region of interest (a circle with 18-24 pixel diameter) with continuous 405 nm or 488 nm lasers controlled by the SIM scanner of the Olympus FV12000 confocal microscope while scanning. The imaging laser had a similar intensity than the stimulus laser but covered an area that was 140-250 times larger than the stimulus ROI.

### Calcium-imaging data analysis and image registration

The calcium-imaging movies were analyzed with FIJI (Schindelin et al., 2012) (RRID:SCR_002285) and a custom Python script, as described previously (Gühmann et al., 2015), with the following modifications. The movies were motion-corrected in FIJI with moco (Dubbs et al., 2016) and Descriptor based registration (https://github.com/fiji/Descriptor_based_registration). The data are shown as ΔF/F0. For the calculation of the normalized ΔF/F0 with a time-dependent baseline, F0 was set as the average of an area of the brain with no calcium activity to normalize for the additional lasers (405 nm and 488 nm) and potential artefacts from the microscope’s detector. Correlation analyses were done using FIJI and Python. The ROI was defined manually and was correlated with every pixel of the time-series. Finally, a single image was created with the Pearson correlation coefficients and a [−1, 1] heatmap plot with two colors.

### Vertical column setup for measuring photoresponses

Photoresponses of larvae of different ages were assayed in a vertical Plexiglas column

(31 mm x 10 mm x 160 mm water height). The column was illuminated from above with light from a monochromator (Polychrome II, Till Photonics). The monochromator was controlled by AxioVision 4.8.2.0 (Carl Zeiss MicroImaging GmbH) via analog voltage or by a custom written Java program via the serial port. The light passed a collimator lens (LAG-65.0-53.0-C with MgF2 Coating, CVI Melles Griot). The column was illuminated from both sides with light-emitting diodes (LEDs). The LEDs on each side were grouped into two strips. One strip contained UV (395 nm) LEDs (SMB1W-395, Roithner Lasertechnik) and the other infrared (810 nm) LEDs (SMB1W-810NR-I, Roithner Lasertechnik). The UV LEDs were run at 4 V to stimulate the larvae in the column from the side. The infrared LEDs were run at 8 V (overvoltage) to illuminate the larvae for the camera (DMK 22BUC03, The Imaging Source), which recorded videos at 15 frames per second and was controlled by IC Capture (The Imaging Source).

### Non-directional UV-light assay

27-hour-old, 36-hour-old, 2-day-old, and 3-day-old *Platynereis dumerilii* larvae were stimulated in the column with UV (395 nm) light from the LEDs on the side. Afterwards the larvae were stimulated with monochromatic blue (480 nm) light coming from above to assay for phototaxis. Each stimulus lasted 4 min. The LEDs were controlled manually and the monochromator (Polychrome II, Till Photonics) was controlled via AxioVision.

### Comparing behavior of wildtype and *c-opsin1* ‐knockout larvae

To compare the behavior of wildtype and *c-opsin1-*knockout larvae in the vertical column, for each genotype we mixed several batches of larvae. Larvae were dark adapted for 5 min. The larvae were recorded for 1 min in the dark followed by exposure to collimated blue (480 nm) light from the top of the column for 2 min, then 2 min darkness, and finally collimated UV (395 nm) light from above for 2 min. Stimulus light was provided by a monochromator (Polychrome II, Till Photonics).

### UV-green-light switching assay

Early and late-stage *Platynereis dumerilii* larvae were assayed in the vertical columns. The larvae were stimulated 6 times alternatively with UV (380 nm) and green (520 nm) light. Each stimulus lasted 4.5 min. The last 3 min within each stimulus were analyzed for vertical displacement of the larvae. The light was provided by a monochromator (Polychrome II, Till Photonics), which was controlled by AxioVision.

### Action spectrum of vertical swimming

2-day-old and 3-day-old *Platynereis dumerilii* larvae were assayed in the vertical columns. The larvae were stimulated with monochromatic light from above between 340 nm and 480 nm in 20 nm steps. Between the stimuli, additional 520 nm stimuli were introduced to avoid the accumulation of the larvae at the bottom after UV treatment. The larvae were also stimulated with monochromatic light from above between 400 nm and 680 nm in 20 nm steps. Between these stimuli, additional 400 nm stimuli were introduced to avoid the accumulation of the larvae at the top due to phototaxis. Each stimulus lasted 3.5 min. The last 2 min of each stimulus were analyzed for vertical displacement of the larvae. The light was provided by a monochromator (Polychrome II, Till Photonics), which was controlled by AxioVision.

### Ratio-metric assay

3-day-old *Platynereis dumerilii* larvae were stimulated with UV-blue (380 nm, 480 nm) or UV-red (380 nm, 660 nm) mixed light from above. The larvae were mixed to be uniformly distributed in the column and dark adapted for 5 min. In each step, 10 % UV-light was replaced by blue or red light. Each step was followed by a blue (480 nm) light stimulus to avoid the accumulation of the larvae at the bottom after UV treatment. Different ratios were created by rapidly switching between wavelengths within 500 ms periods (e.g., for a 10% UV 90% blue ratio we provided UV-light for 50 ms followed by blue light for 450 ms). Each stimulus condition lasted 4 min. The monochromator was controlled by a custom Java program via the serial port.

### Vertical cuvette photoresponse assay

2-day-old *Platynereis dumerilii* larvae were assayed in a vertical cuvette of 10 mm x

10 mm x 42 mm (L x W x H). The larvae were kept at 18°C and had been exposed to daylight before the experiment. The larvae were illuminated with a monochromator (Polychrome II, Till Photonics) from one side of the cuvette. A mirror (PFSQ20-03-F01, Thorlabs) placed at the opposite side reflected the light to provide near-uniform bilateral illumination. The light passed a diffuser (Ø1” 20° Circle Pattern Diffuser, ED1-C20; Thorlabs) and a collimating lens (LAG-65.0-53.0-C with MgF2 Coating, CVI Melles Griot) before it hit the cuvette. The cuvette was illuminated with infrared (850 nm) light-emitting diodes (LEDs) (L2X2-I3CA, Roithner Lasertechnik) from the side. The LEDs were run at 6.0 V. The larvae were stimulated with light from 340 nm to 560 nm in 20 nm steps. Each step lasted 1 min. The steps were separated by 1 min darkness, so that the larvae could redistribute after each stimulus. The larvae were recorded at 16 frames per second with a DMK 23GP031 camera (The Imaging Source) controlled by IC Capture. The camera was equipped with a macro objective (Macro Zoomatar 1:4/50-125 mm, Zoomar Muenchen). It was mounted with a closeup lens (+2 52 mm, Dörr close-up lens set 368052). The larvae were tracked and their vertical displacement was analyzed during the last 45 s of each stimulus period. Scripts are available at https://github.com/JekelyLab/Depth_gauge.

## Acknowledgements

The research leading to these results received funding from the European Research Council under the European Union’s Seventh Framework Programme (FP7/2007-2013)/European Research Council Grant Agreement 260821. The research was supported by a grant from the DFG - Deutsche Forschungsgemeinschaft (Reference no. JE 777/3-1). S. Y. was supported by the National Institutes of Health (R01EY016400) and Emory University.

---

**Supplementary Figure 1.**
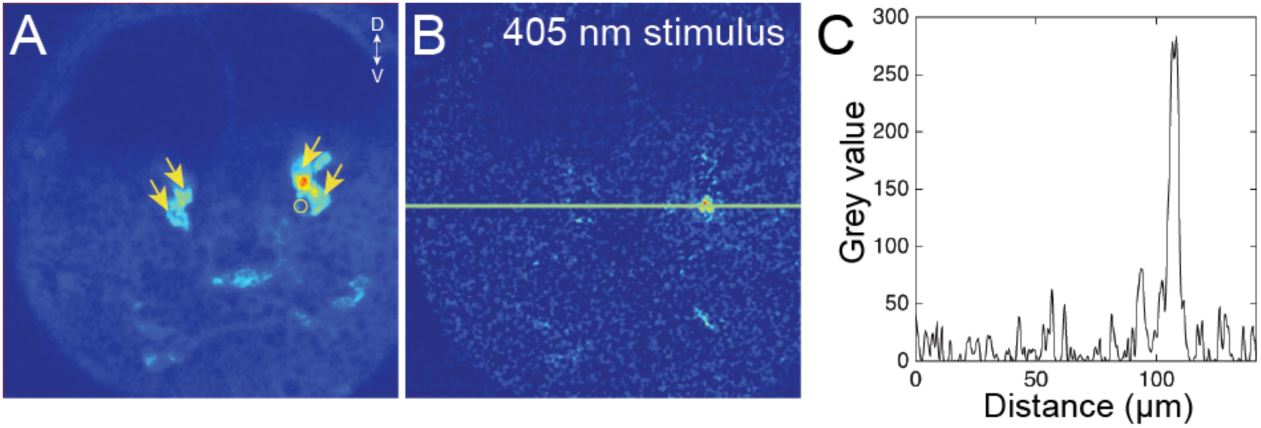
Quantification of stimulus light intensity during the local stimulation of cPRC cilia. (A) The four cPRCs (arrows) are recognized based on their high GCaMP6s fluorescence. A region of interest used to deliver the 405 nm stimulation with the independent SIM scanner is shown (circle). (B) Distribution of 405 nm light scattered by the sample. (C) Quantification of signal intensity based on the line shown in (B).

**Supplementary Figure 2.**
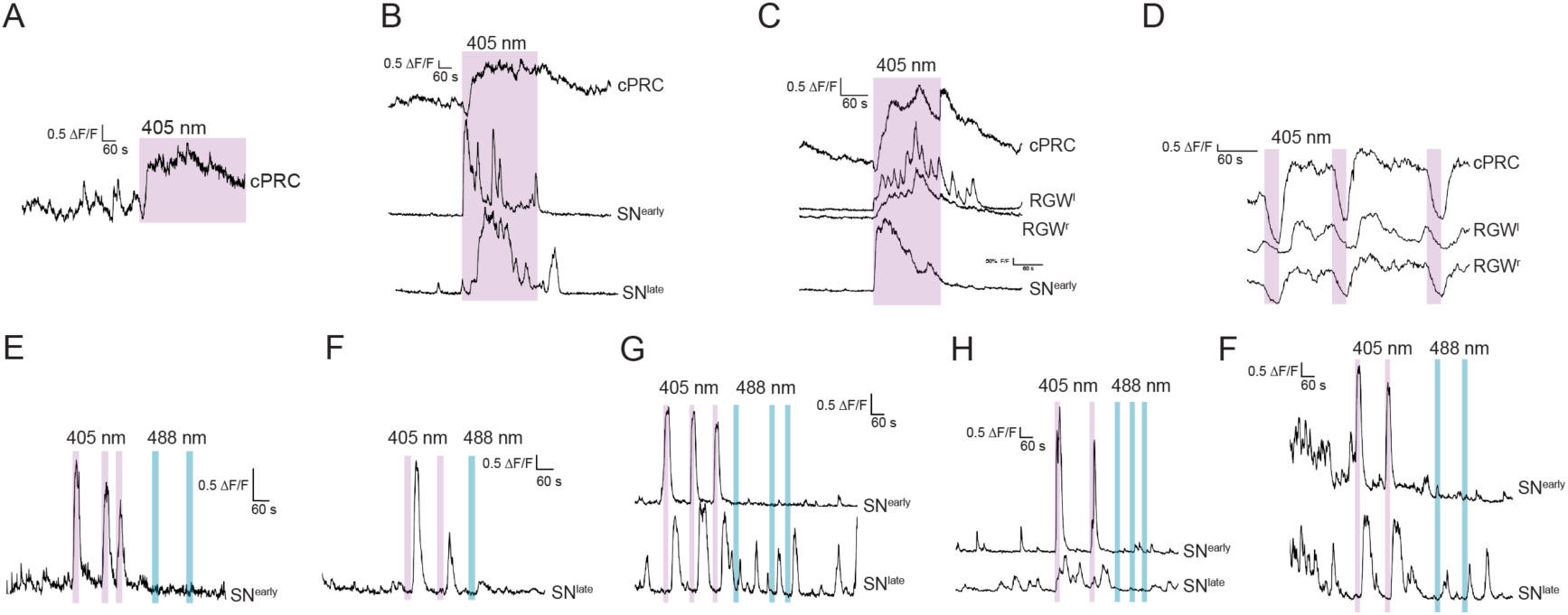
Calcium imaging in *Platynereis* larvae combined with the stimulation of cPRCs. (A) cPRC response to prolonged local 405 nm stimulation. (B) Response of a cPRC, an SN^early^ and an SN^late^ sensory neuron to prolonged 405 nm stimulation. (C) Response of a cPRC, two RGW interneurons, and an SN^early^ sensory neuron to prolonged 405 nm stimulation. (D) Responses of a cPRC and two RGW interneurons to repeated 405 nm stimulation of 20 sec duration. (E) Responses of an SN^early^ sensory neuron to repeated 405 nm and 488 nm stimulation of 20 sec duration. (F) Responses of an SN^late^ sensory neuron to repeated 405 nm and 488 nm stimulation of 20 sec duration. (G-F) Responses of SN^early^ and SN^late^ sensory neurons to repeated 405 nm and 488 nm stimulation of 20 sec duration.

Movie 1. Wiring diagram of cPRC and rPRC circuits in the *Platynereis* larval head. The anatomy of the reconstructed neurons is shown together with the position of the same cells in the wiring diagram.

Movie 2. Behavioral responses of *Platynereis* larvae to 380 nm illumination from above. The magenta square in the top corner marks when the UV stimulus light was switched on. The tracks are color-coded based on heading direction (red, upward; blue, downward). The movie is sped up 2x.

